# Dual Role of Glycosylation in Resistance to CD4-binding Site Broadly Neutralizing Antibodies

**DOI:** 10.1101/2025.07.15.664896

**Authors:** Teresa Murphy, Meagan Kelly, Kai S. Shimagaki, Thomas DeStefanis, Gabriel Galeotos, Myungjin Lee, Qing Wei, Jan Novak, Katharine J. Bar, John P. Barton, Rebecca M. Lynch

**Affiliations:** Department of Microbiology, Immunology and Tropical Medicine, George Washington University; Department of Computational and Systems Biology, and Department of Physics and Astronomy, University of Pittsburgh; Vaccine Research Center, National Institute of Allergy and Infectious Diseases, National Institutes of Health, Bethesda, MD 20892, USA; Department of Microbiology, University of Alabama at Birmingham; Department of Medicine, Perelman School of Medicine, University of Pennsylvania, Philadelphia, Pennsylvania, USA

**Keywords:** Broadly neutralizing antibodies, HIV escape, glycosylation, CD4 binding-site antibodies

## Abstract

Broadly neutralizing antibodies (bNAbs) provide a useful tool for HIV cure strategies because of their ability to target conserved regions on the envelope (Env) protein in the context of both virions and infected cells. One of the most well studied bNAbs is the CD4 binding site (CD4bs) antibody, VRC01 and others in its class. A major obstacle to effective cure strategies with bNAbs is viral immune escape. A deeper understanding of escape pathways from VRC01-class antibodies in genetically diverse samples is needed. Using an *in vitro* viral escape assay where infected CD4^+^ T cells were cultured in the presence of increasing VRC01 concentrations, complete resistance to VRC01 was detected in Env 246.F3 by day 42. We determined that resistance was due to a mutation at position N276 that resulted in elimination of the glycan-attachment site. As the loss of a glycan at this site is known to increase virus sensitivity to CD4bs antibodies, we explored this finding further. Specifically, we introduced N276D mutation into the 12-virus global Env panel and measured neutralization susceptibility to a panel of CD4bs bNAbs. This N276D mutation increased resistance or sensitivity depending on the Env and bNAb in question, emphasizing a dual role of glycan N276 in VRC01-class bNAb neutralization. The role of this glycan in escape was demonstrated to be dependent on the context of the Env, the bNAb and the glycosylation complexity of the virus due to producer cells. These findings underscore the complexity of glycosylation in genetically diverse HIV escape from antibodies.

**Importance:** Despite decades of research, an HIV cure remains elusive, largely due to the virus’s immense genetic variability and ability to evade immune clearance. While antiretroviral therapy (ART) suppresses viral replication, it does not eradicate the virus and presents long-term challenges related to toxicity, access, and adherence. Broadly neutralizing antibodies (bNAbs), particularly those targeting the conserved CD4 binding site such as VRC01 and others in its class, offer promise for durable control or cure by targeting both circulating virus and infected cells. However, viral escape from bNAbs remains a critical hurdle. In this study, we demonstrate that glycan-mediated escape from VRC01-class bNAbs is highly context-dependent—shaped by Env, bNAb, and the glycosylation patterns introduced by the producer cell. These findings emphasize the dual role of glycans in affecting antibody sensitivity and underscore the importance of viral and host factors in shaping effective bNAb-based cure strategies across diverse HIV-1 strains.

## Introduction

The HIV envelope glycoprotein (Env), which is a trimer of gp120 and gp41 dimers on the surface viral particles, is the dominant target for anti-HIV neutralizing antibodies, including broadly neutralizing antibodies (bNAbs)^1,2,3^. Because of high levels of replication and an error-prone polymerase, there are many mechanisms by which HIV’s Env protein evades this neutralization and clearance by antibodies, including structural occlusion of conserved epitopes, shifting glycosylation sites and high sequence variation ^4,5^. Structural occlusion is the use of post-translational modifications, such as glycans, to protect the viral envelope from being bound by immune cells. N-linked glycans are post-translational modifications on the Env protein that not only protect the virion from neutralization by shielding the epitopes targeted by antibodies, but also aid in protein folding and viral infectivity ^5–7^ and are mainly invisible to the host’s immune response as they are part of the host post translational machinery and therefore are “self”. Although these glycans mainly shield neutralizing epitopes, they can also comprise portions of the epitopes targeted by certain bNAbs^8^. One such site that heavily utilizes this protective glycosylation pattern is the CD4-binding site (CD4bs). This highly conserved site is the region of Env responsible for virion cell entry by binding to the receptor CD4 on immune cells to mediate infection. The CD4bs is not only hidden structurally in a pocket, but also surrounded by 4 main glycans: positions N197, N276, N363 and N462 ^9^. These residues are conserved because they comprise the crucial viral function of receptor binding, which means that bNAbs targeting this site are fairly broad and potent as they bind to many of the same residues on the Env’s surface that CD4 itself binds to ^10,11^. Many CD4bs-targeting bNAbs have been isolated to date, of which several have similar genetic properties to one of the original bNAbs, VRC01, and are referred to as VRC01-class^12^. These CD4bs-targeting bNAbs have been well characterized functionally and structurally^10^.

Although this region is surrounded by 4 highly conserved glycans, their role in antibody neutralization can be confusing because they both can increase resistance to certain neutralizing antibodies through shielding while also serving as part of the neutralizing epitope for other antibodies. Examples can be observed in viral escape occurring in individuals who generated these bNAbs. In the viruses from the VRC01 donor, the addition of a glycan at position 276 conferred a 10-fold increase in the IC50 of VRC01^3^, indicating its role in escape from this antibody response. This role of this glycan in shielding the CD4bs is reinforced by studies demonstrating that removal of glycan 276 increased neutralization sensitivity to VRC01-like bNAbs ^9^ or binding to gp120 ^13^. Furthermore, when designing vaccine immunogens, the glycans surrounding the CD4bs, including N276, are commonly removed in order to increase exposure of this site to naïve B cells ^9,14^. However, this glycan is highly conserved, and so the ability to accommodate this glycan leads to greater bNAb neutralization breadth, as demonstrated by studies where bNAbs are reverted to germline (naïve-like BCR), and lose their ability to neutralize viruses with glycan 276 ^15,14^. Furthermore, in contrast to VRC01-class bNAbs, there is another class of CD4bs bNAbs, such as HJ16 and 179NC75, for which the N276 glycan and its removal confers complete neutralization resistance, ^16–18^. In the individual from whom 179NC75 was isolated, loss of glycan 276 was identified as an escape mutation from 179NC75 ^17^, and this glycan’s loss has been identified as an escape mutation from autologous antibodies in subtype C infection as well^13^. This effect is due to the direct contact bNAb 179NC75 heavy chain makes with the N276 glycan ^19^. Thus, the dual role of glycan 276 in CD4bs bNAb epitopes renders this glycan neither a universal resistance or sensitivity signature for CD4bs bNAbs ^20^.

The impact of this glycan in both vaccine design and in viral escape from bNAbs emphasizes the need for a deeper understanding of its role in the neutralization profiles of CD4bs bNAbs. Much of the *in vitro* work performed to date examines the effects of glycan removal in subtype B Envs, such as YU2, or in the context of vaccine design, which has focused on subtype A Env BG505 ^7–12^ and subtype C Env 426c. This lack of genetically diverse *in vitro* research leads to gaps in knowledge before bNAb clinical trials are initiated. VRC01 has been included in a multitude of clinical trials that have ranged across various geographic areas, in viremic individuals and those suppressed on ART as well as prevention trials with the issue of viral resistance always arising by the end of the trial ^21^. A more in depth understanding of viral escape from bNAbs using *in vitro* methods will allow study of the impacts of glycan removal across genetically diverse HIV subtypes, which is important for future bNAb clinical trials

To study how non-subtype B viruses escape from VRC01, we performed an *in vitro* viral escape assay with the subtype AC Env 246.F3. We observed that complete neutralization escape was mediated by a mutation of N276K, which also removes a potential N-linked glycosylation at this position. Therefore, we investigated the effects of this mutation on neutralization susceptibility across genetically diverse subtypes of HIV-1. Our analysis indicated that the initial resistance to VRC01 was most likely conferred by the residue change at position 276 but that each VRC01-class bNAb was differently affected. These findings highlight the complex role of glycan shielding in bNAb neutralization resistance and heterogeneity even between bNAbs that are genetically and structurally similar and target the same site on the virus.

## Methods

### HIV DNA plasmids

The infectious molecular clone (IMC) 246.F3-NL4.3+BN was derived by subcloning the subtype AC 246.F3 *env* gene into a replication competent NL4.3 backbone (NIH AIDS Repository) while additionally adding in BstEII and NcoI restriction sites using a previously described cloning strategy^22^. Briefly, the NcoI and BstEII restriction sites were inserted into pNL4.3 plasmid by GenScript (Piscataway, NJ) so that they flank the *env* gene, leaving 37 AA at the 5’ and 8 at the 3’ end of the Env as wildtype NL4.3 although virtually the entire desired *env* gene is present. The 246.F3 *env* plasmid (BEI Resources, NIH HIV Reagent Program) was PCR amplified with previously described primers C-CstEII and TNE3*-*NcoI to introduce flanking BstEII and NcoI restriction sites around the envelope with Phusion HF polymerase (ThermoFischer Scientific, Waltham, MA). PCR product was PCR purified with the QIAquick PCR Purification Kit (Qiagen, Hilden, DE). Double digestion with NcoI and BstEII enzymes was performed on 30 μL of purified PCR product for the *env* and 5 μL of NL4.3+BN. In order to increase ligation efficiency, the NL4.3+BN backbone was dephosphorylated with Antarctic Phosphatase at 37°C. Ligation was performed on digested envelope and backbone products with the Quick Ligation Kit (New England Biolabs, Ipswich, MA). Plasmids were transformed using XL-10 Gold competent cells and plated on LB-ampicillin plates. After 24 hours, bacterial colonies were picked and grown in LB-Amp broth for 24 hours before being mini-prepped with the QiaPrep Spin MINIPrep kit (Qiagen, Hilden, DE). Plasmid *envs* were sequence verified.

Wildtype *env* gene plasmids from the global panel of HIV-1 reference strains were obtained from the NIH HIV Reagent Program. Plasmids containing the N276D mutation were generated via site-directed mutagenesis as follows. Completely overlapping primers between 40-50 bps designed to insert the N276D mutation into each *env* were synthesized (IDT) and used to amplify each plasmid according to the QuikChange Lighting Site-Directed Mutagenesis kit (Agilent, Santa Clara, CA) instructions. Amplicons were transformed in Stbl2 competent cells and plated on LB-ampicillin plates and harvested after 24 hours at 30°C, and DNA was isolated and verified as described for the IMC.

### Generation of virus stocks

Wild type and mutant pseudovirus stocks were generated by co-transfecting 293T cells with the env plasmid and an env-deficient backbone (pSG3 Δenv) at a 1:3 ratio by mass of DNA while IMC stocks were transfected with 13.3 μg of DNA. To test differential glycosylation patterns, pseudoviruses were generated in Expi293 cells as follows. Cells were seeded at 2.0×10^6^ viable cells/ml 24 hours before transfecting 20 μg of env-deficient backbone (pSG3 Δenv) and 10 μg of desired env plasmid. For all virus stock, culture supernatants were collected 72 hours after transfection, and were harvested, filtered, aliquoted and frozen at −80°C until further use.

### *In Vitro* bNAb Resistance Assay

Uninfected, target cells were isolated from buffy coat obtained from GulfCoast Blood Bank (Houston, TX) by isolating PBMC using SepMate PBMC Isolation tubes (STEMCell, Vancouver, BC) and CD8-depleting these cells with CD8^+^ Dynabeads (ThermoFischer Scientific, Waltham, MA) according to the manufacturer’s instructions. These CD4^+^ enriched PBMC were cultured in complete RPMI medium for 3 days in the presence of 20 g/ml phytohemagglutinin (PHA) for activation prior to infection. Virus stock of 246.F3-NL4.3+BN at an MOI of 1 was incubated with bNAb VRC01 at two concentrations: 0.35 μg/ml and 0.85 μg/ml, for 30 minutes. 100 μl of uninfected CD4^+^ enriched PBMC at 1×10^6^ cells/ml were added to each infection and incubated for 2 hours in a low volume incubation before being supplemented to 2 ml with complete RPMI medium supplemented with 20 U/ml recombinant human interleukin-2 (IL-2) (Roche Diagnostics). After 24 hours, all infected cells were plated in 12 well plates and cultured for up to 42 days in the presence of increasing concentrations of VRC01. Every 2 to 3 days, half of the supernatant was refreshed with new IL-2 media and VRC01. 200 μL of old supernatant was frozen for p24 analysis with the AlphaLISA HIV p24 Biotin-Free detection kit (Revvity, Waltham, MA). Every 14 days, target cells were replenished by spinoculating freshly activated *ex vivo* CD4^+^ T with cell-free viral supernatant for each well. Remaining CD4^+^ T cells that had been in culture for the previous 14 days were replenished with antibody-free complete RPMI medium for 24 hours. Aliquots of viral supernatant were then frozen for neutralization assays and viral sequencing. Schematic of the experiment design is shown in Fig. 1.

**Figure 1.**
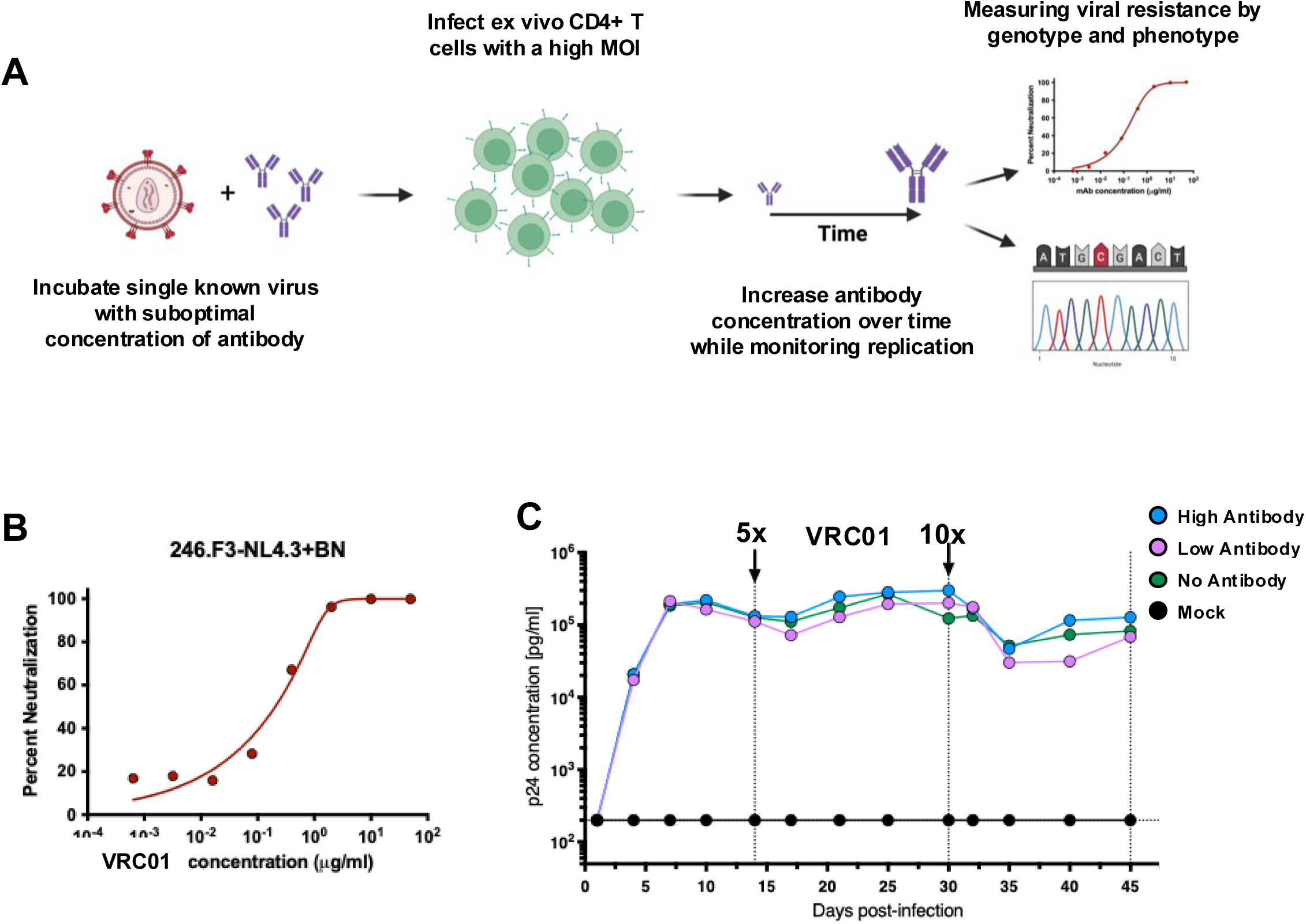
Schematic of 246.F3-NL4.2+BN *in vitro* viral resistance assay. A. Virus and bNAb pairing are incubated for 30min. before a high MOI standing infection of pre-stimulated CD4 enriched ex vivo T cells. Infections are plated and monitored for replication kinetics by measuring p24 every 3 days. Every 14 days, supernatant is used to infect new CD4 T cells for infection propagation. New media without any bNAb is added to infected cells for virus to grow for 24 hours. This cell free/antibody free virus is then measured for resistance phenotypically and genotypically. B. Stock sensitivity of 246.F3-NL4.3+BN to VRC01 in TZM-bl cells. C. Replication kinetics of p24 concentration was measured for 45 days. Dotted lines indicated when cultures were refreshed with ex vivo CD4+ target cells. Arrows indicate increases in antibody concentration.

### Sequencing viruses using single genome sequencing (SGS)

Sequences were obtained by SGS as previously described ^23^. Briefly, viral RNA was extracted from 140 μl of each well’s virus supernatant by QIAmp kit (Qiagen, Germantown MD). cDNA was synthesized and *env* genes were amplified by nested PCR using the Platinum Taq High Fidelity polymerase (Invitrogen). Template cDNA was serially diluted so that fewer than 33% of PCR replicates were positive, ensuring that a majority of amplicons would be generated from a single cDNA template, according to Poisson distribution. Well-described primers Env_outF1 and Env_outR1 were used for first round amplification, and Env_inF2 and Env_inR2 for the second round. All PCR mixes were generated in PCR clean rooms free of post-PCR or plasmid DNA. Amplicons were run on 1% agarose gels and sequenced by ACGT Inc. A minimum of five single-template sequences were obtained from each well. Sequences that contained stop codons, large deletions, or mixed bases were removed from further analysis.

### Identification of resistance mutations

All confirmed sequences were translated and aligned by MUSCLE to the virus stock Env sequence, which was set as the reference. Amino acid highlighter plots were generated using the Los Alamos National Laboratories Highlighter tool by comparing experimental sequences to the sequence of the infecting 264.F3+NL4.3+BN strain ^24^. Amino acid mutations observed in more than half of an experimental condition were considered to be fixed and further analyzed.

### TZM-bl neutralization assay

This assay was run as previously described ^25–27^. Briefly, input virus dilution of pseudovirus and IMC stocks were calculated from titration experiments to ensure sufficient luciferase output within the linear portion of the titration curve (45,000 RLUs). Culture supernatant from the resistance assay was tested undiluted. All replication competent viruses were run in the presence of 1 μM indinavir to prevent viral replication. 10 μl of five-fold serially diluted mAbs from a starting concentration of 50 μg/ml were incubated with 40 μl of virus in duplicate for 30 minutes at 37°C in 96-well clear flat-bottom black culture plates (Greiner Bio-One). TZM-bl cells were added at a concentration of 10,000 cells per 20 μl to each well in DMEM containing 75 μg/ml DEAE-dextran Cell only and virus only controls were included on each plate. Plates were incubated for 24 hours at 37°C in a 5% CO_2_ incubator, after which the volume of culture medium was adjusted to 200 μl by adding complete DMEM. 48 hours post-infection, 100 μl was removed from each well and 100 μl of SpectraMax Glo Steady-Luc reporter assay (Molecular Devices, LLC., CA) reagent was added to the cells. After a 10-min incubation at room temperature to allow cell lysis, the luminescence intensity was measured using a SpectraMax i3x multi-mode detection platform per the manufacturers’ instructions. Neutralization curves were calculated by averaging duplicate wells and comparing luciferase units of wells containing antibody to virus-only controls after background subtraction. Curves are fit by nonlinear regression using the asymmetric five-parameter logistic equation in Prism 9 for macOS (GraphPad Software, LLC). The 50% and 80% inhibitory concentrations (IC_50_ and IC_80_) are estimates of the antibody concentrations required to inhibit infection by 50% and 80%, respectively.

### Generation of broadly neutralizing antibodies

The heavy- and light-chain genes of VRC01, VRC07-523, VRC13, 3BNC117, N6, 10-1074, PGDM1400 or 10E8v4-5R+100cF were expressed as full-length IgG1s by transient transfection of 293F cells. Protein was purified by affinity chromatography using HiTrap Protein A HP Columns (GE Healthcare).

### Mutational cost analysis

The mutational cost of individual substitutions was estimated using a pairwise Potts model trained on a diverse set of HIV-1 Env sequences ^28–30^. This model captures both the statistical patterns of the sequence ensemble and the underlying fitness landscape. Mutational cost is defined as the energy difference between the mutant and reference sequences under the Potts energy function. Similar approaches were employed and successfully captured the underlying fitness effects of individual mutations ^31–33^. Details of the training data and model are provided below.

#### Training data

HIV-1 Env amino acid sequences from all subtypes were retrieved from the HIV sequence database at Los Alamos National Laboratory ^34^ and aligned to the HXB2 reference sequence using HIValign ^35^, resulting in over 100,000 sequences. The sequence ensemble is globally well-aligned, and we restricted our analysis to sites that do not correspond to alignment gaps in the HXB2 sequence, yielding a final sequence length of 856 amino acids. Additionally, to control for data quality, we excluded sequences with more than 20% alignment gaps relative to the total sequence length, resulting in 92,388 sequences derived from 8,386 unique subjects. Naively treating individual sequences equally, introduces bias in the statistical analysis, as the number of sequences per subject varies widely, some subjects contribute over a thousand sequences, while most contribute only a handful. To correct for this imbalance, we estimated per-sequence weights inversely proportional to the number of sequences from the same individual (i.e., weight *∝* 1/*n*, where *n* is the number of sequences from a given subject).

#### Model

The pairwise Potts model, formulated as a Gibbs distribution with a pairwise energy function ^28–30^, was trained to reproduce essential statistical features of the sequence ensemble, such as the site-specific amino acid frequencies and the pairwise covariances across sites. Training the model involves optimizing the parameters of the energy function using maximum likelihood estimation. Mathematically, let the amino acid sequence be denoted by *A* = (*A*_1_,…, *A_L_*), where *L* is the sequence length (*L* = 856 in this study). The energy function is defined as *E*(*A* ∣ *h*, *J*), where *h* and *J* are model parameters to be learned. The corresponding probability distribution over sequences is: 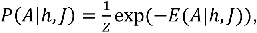 where *Z* is the normalization constant (partition function).

The explicit form of the energy function is:

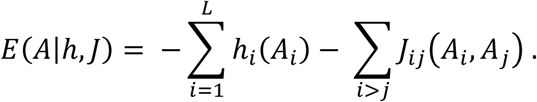

Here, *h_i_*(*a*) adjusts the frequency of amino acid *a* at site *i*, while *J_ij_*(*a*, *b*) captures pairwise interactions between amino acids *a* and *b* at sites *i* and *j*, respectively. The pairwise parameters *J* encode epistatic interactions, which can reflect co-evolution of residues and underlying functional or structural constraints ^28–30^. Once the parameters *h* and *J* are optimized, the energy function serves as a proxy for the fitness landscape. The mutational cost of substituting an amino acid at site *i* in a reference sequence *A*^∗^ (e.g., the stock strain 246.F3-NL4.3+BN, in this study) with an amino acid *a* is computed as:

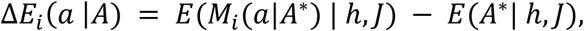

where *M_i_*(*a* ∣ *A*) denotes the sequence obtained by replacing the amino acid at position *i* in *A* with *a*: *M_i_*(*a* ∣ *A*) = (*A*_1_,…, *A_i_*_-1_, *a*, *A_i_*_+1_,…, *A_L_*).

#### Training details

In principle, model parameters are learned by maximizing the likelihood, which is equivalent to matching expected statistics *O* (e.g., site-specific frequencies and pairwise covariances) between the data and the model:

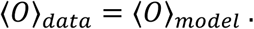

Here, ⍰*O*⍰*_data_* = ∑*_m_ W_m_O*(*A^m^*) / ∑*_m_ W_m_*, ⍰*O*⍰*_model_* = ∑*_A_ O*(*A*)*P*(*A*|*h*, *J*), and *W_m_* denotes the weight of sequence *A^m^*, as previously defined. Naive optimization is computationally demanding due to the intractability of the partition function *Z*, which sums over exponentially many sequences. While Monte Carlo sampling can be used ^31^, several more efficient models have been proposed for efficient training ^28–30,36–40^. In this study, we employed the pseudo-likelihood maximization approach, which assumes conditional independence of sites given the rest of the sequence ^30^. This method efficiently estimates both *h* and *J*, and has been widely adopted for large-scale Potts model inference.

### Homology modeling of HIV-1 Env-VRC01 complexes

Homology modeling was conducted using YASARA Structure (www.yasara.org). A total of 12 HIV Env-VRC01 complex structures were modeled using Env sequences from a global panel of 12 viruses. The BG505-VRC01 structure (PDB: 6V8X) was used as the single structural template. The target and template sequences were aligned using MAFFT (Katoh et al, Nucleic Acids Research, 2002) prior to modeling. The modeling process followed YASARA’s standard homology modeling pipeline, which includes optimization and refinement using steepest descent and simulated annealing energy minimization.

## Results

### Identification of subtype AC 246.F3 envelope escape mutations from VRC01 *in vitro*

In order to study escape from bNAb VRC01 in a non-subtype B virus, we generated an infectious molecular clone (IMC) with the subtype AC 246.F3 envelope inserted into an NL4.3 backbone. We employed an *in vitro* viral escape assay in which the concentration CD4bs bNAb VRC01 was increased over time to allow for selection of mutations that confer resistance (Fig 1A). In this specific experiment, the two initial VRC01 concentrations that were chosen based on the virus stock sensitivity to the VRC01 (Fig 1B). The IC_50_ and IC_80_ were 0.35 μg/ml, referred to as low antibody, and 0.85 μg/ml, referred to as high antibody, respectively. Resistance was determined phenotypically and genotypically every 14 days during the assay. Between days 30 and 45, as the antibody concentration was increased from 4.2 μg/ml to 42 μg/ml for the high antibody concentration and 1.75 μg/ml to 17.5 μg/ml in the low antibody concentration, complete neutralization resistance developed in the two wells where virus was replicating in the presence of antibody (in both high and low concentration conditions (Fig 2A)). Envs from all three wells (no, high and low antibody) were sequenced and aligned to the stock Env as a reference. Four mutations in the alignment were observed compared to the reference. Two of these were only observed in the no antibody condition; one was only found in the low antibody condition and one was observed in both antibody conditions. Of the two fixed mutations found in the antibody conditions, one was L122I in the gp120 protein before the V2 loop and only observed in the low antibody well, while the second was N276K in Loop D, and was observed in both the low and high antibody wells (Fig 2B). These same sequences were additionally analyzed for number of mutations and the cost of each mutation in terms of Pott’s Energy compared to the no antibody controls. Overall, the cost of the fixed mutations found in the *in vitro* resistance experiment were generally within lower Pott’s Energy readings, corresponding with lower or neutral mutation costs (Fig 2C). Specifically, the mutations observed in the no antibody control wells have opposing fitness costs, with V181I conferring a slight fitness benefit and S189R having a slight fitness cost, implying that there may be a synergistic relationship or a linkage between these two mutations, as they are only seen in tandem with one another in the no antibody control wells. Indeed, the epistatic interaction between these mutations is negative; that is, co-occurrence is preferred. Thus, the fitness model suggests that the co-occurrence of mutations 181I and 189R can reduce the mutation cost. It is of note that at position 181, isoleucine (I) is heavily conserved in subtype B viruses and valine (V) being more conserved in non-subtype B viruses, with I/V being the two most commonly found residues at this position ^41,42^. The fact that the 2 dominant mutations in the antibody wells (L122I) and (N276K) may exert a fitness cost to the virus, highlights how beneficial they may be in the presence of the antibody VRC01, where the fitness environment in not solely replication but also antibody resistance.

**Figure 2.**
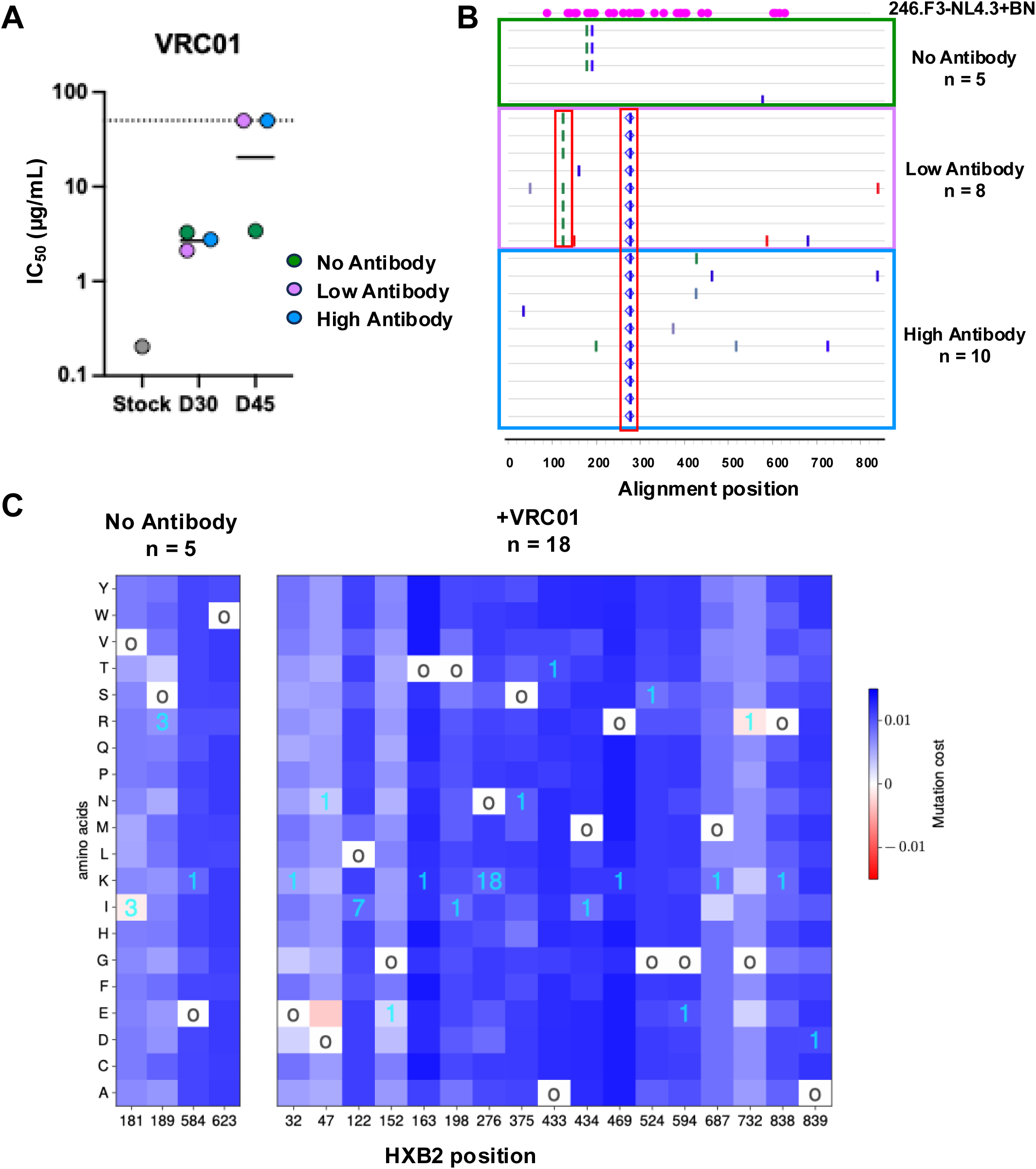
Identification of escape mutations from VRC01. **A**. Antibody-free viral supernatant from 3 wells was tested for sensitivity to VRC01 by TZM-bl neutralization assay at Days 30 and 45 post infection. IC50s were calculated, and an IC50 >50 ug/mL indicates resistance. Geometric mean is plotted. Dotted line indicates the highest bNAb concentration tested (50 ug/mL). **B**. Single genome sequencing was performed to obtain *env* sequences in all three wells at Day 45 and compared to stock virus. Glycans in the stock are indicated in pink. Changes that result in a putative glycan removal are indicated by blue open diamonds. **C**. The x-axis represents amino acid positions (1 to 856) of detected mutations and the y-axis lists all possible residues. Fitness cost of mutations were calculated with Pott’s energy. Lower fitness costs are displayed in red, while higher fitness costs are shown in blue. The color scale is normalized using the maximum and minimum fitness values among all sites. 18 experimental sequences from both antibody conditions and 5 no antibody control sequences were analyzed. The open circles indicate the amino acid at that position in the original virus stock sequence The number of sequences containing the listed mutation are shown in cyan.

### Prevalence of identified VRC01 escape mutation at position 276

In order to confirm individual mutation resistance profiles, single mutations were inserted into the 246.F3 *env* gene expression plasmid and used to generate pseudoviruses. While L122I conferred no neutralization resistance to VRC01, the removal of the N276 glycan via a mutation from asparagine to lysine conferred complete neutralization resistance causing an IC50 change from 0.204 μg/ml to >50 μg/ml (Fig 3A). Over 92,000 genetically diverse *env* genes from Los Alamos National Laboratories’ database were analyzed in order to determine the residue conservation at position 276. Approximately 98% of the sequences have N at position 276, while fewer than 0.2% have K at that position. Other mutations, such as D and S, are also noted at moderate frequencies (Fig 3B). We next analyzed if 276K prevalence varied across subtypes. Out of a total of 253 sequences containing the 276K, it was predominantly observed in subtype C (105 sequences) followed by B (58 sequences) and then CRF01_AE (50 sequences), and minorly found in other subtypes (Fig 3C).

**Figure 3.**
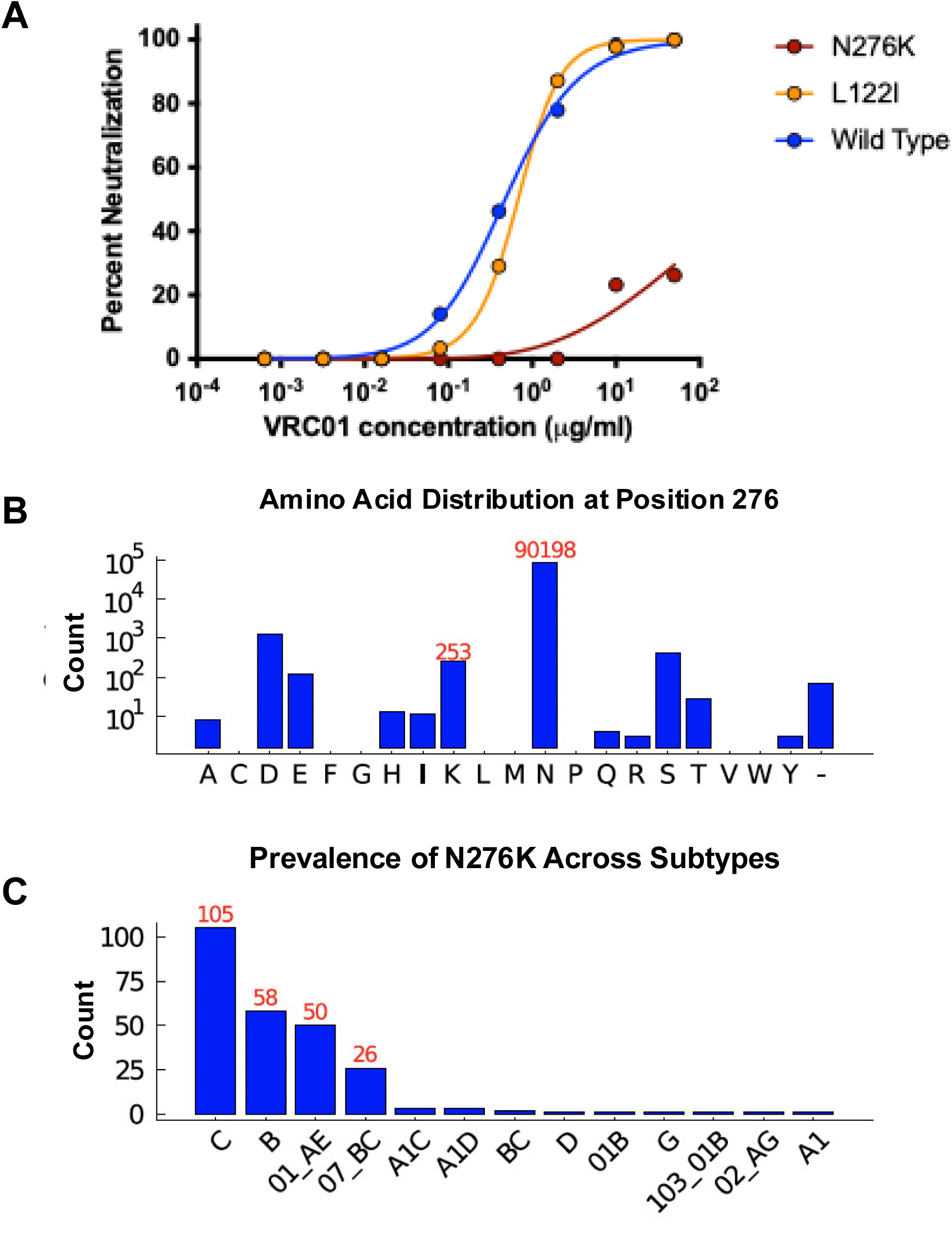
Prevalence of VRC01 escape mutations at 276. **A.** Pseudoviruses of 246.F3-SG3 WT Env with the mutations identified in VRC01 wells were tested for sensitivity to VRC01. **B**. Frequency of sequences with the indicated amino acids at position 276 in the LANL database. **C**. Of 253 sequences containing the K276, the number within each subtype is graphed.

### Effect of N276 glycan removal on neutralization profiles of CD4bs bNAbs

To determine the effect of glycan removal on CD4bs bNAb neutralization in genetically diverse HIV-1 Envs, the N276D mutation was inserted into each of the *env* plasmids from the global panel of HIV-1 reference strains via site-directed mutagenesis. In general, mutation from N to D at 276 increased sensitivity or had no significant change to neutralization susceptibility to 4 CD4bs targeting bNAbs: VRC01, VRC07-523, 3BNC117 and N6 (Fig 4A) when measured as change greater than 5-fold in IC_50_, which was chosen as a conservative increase of the typical 3-fold variation on the pseudovirus neutralization assay. We included antibodies to other sites on the virus (V2 apex PGDM1400, V3-glycan 101-1074 and MPER 10E8v4-V5R+100cF) to control for overall changes to the trimer structure. For the most part, PGDM1400 and 10-1074 neutralization of the virus panel were unaffected by this N276D mutation, but the gp41 MPER targeting antibody did seem to become more potent against certain viruses (Sup Fig 1A). Overall, of the 4 CD4bs-targeting bNAbs, VRC01, VRC07-523 and N6 were all significantly more potent against the N276D virus as measured by Wilcoxon test (Fig 4B). 3BNC117 was more heterogeneous in the resulting neutralization susceptibility (Fig 4B). Control antibodies 10E8v4-5R+100cF and 10-1074 did not demonstrate a statistically significant change in neutralization sensitivity, whereas PGDM1400 did (Sup Fig 1B). Noticeably, subtype B TRO.11-N276D was the only Env to become resistant to VRC01 but not 246.F3-N276D, contradicting our prior results suggesting the loss of glycan at 276 conferred resistance to 246.F3.

**Figure 4.**
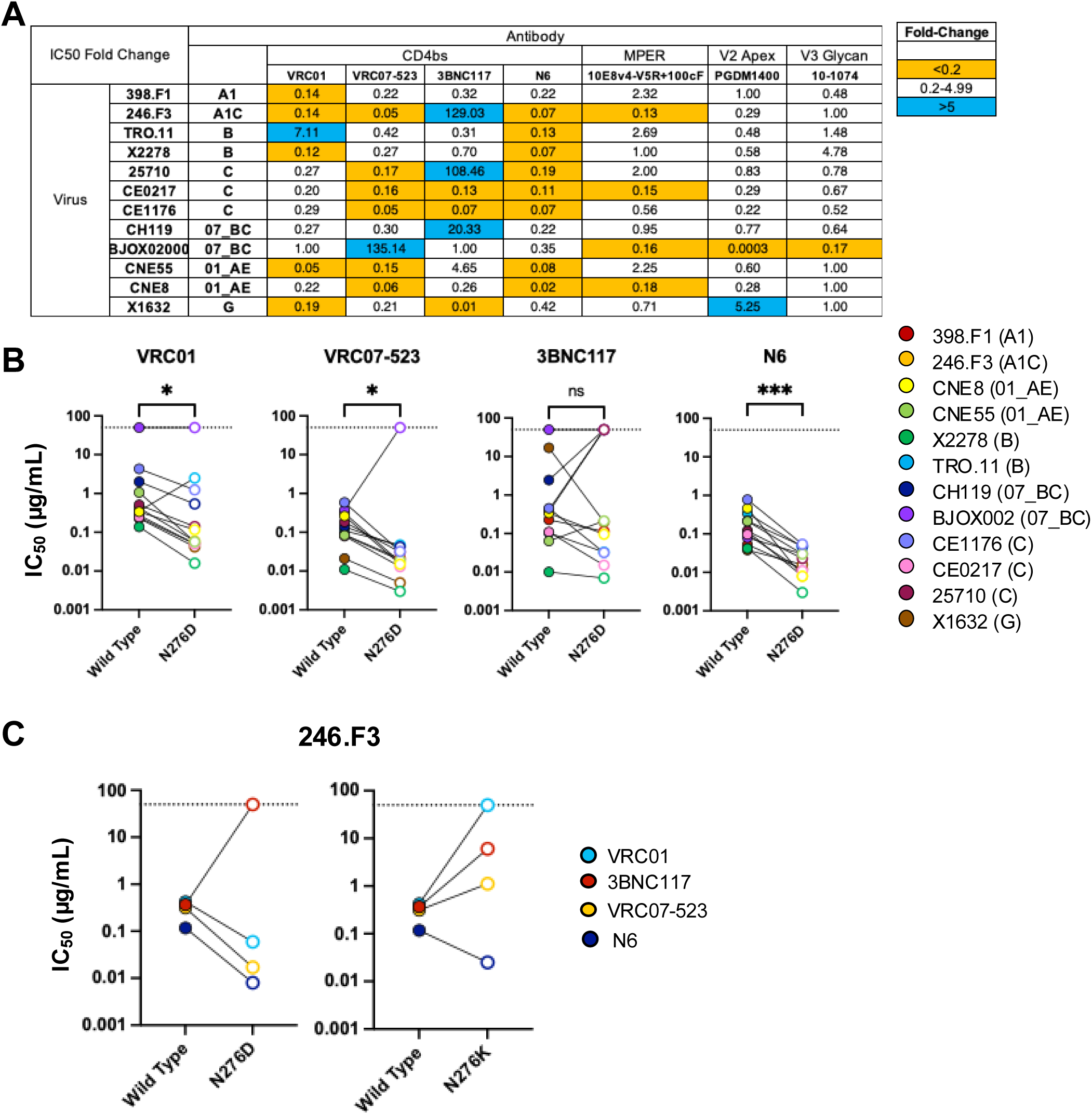
Effect of N276 glycan removal on neutralization profiles of CD4 bNAbs. A. The fold-change of the IC50 of each virus-antibody pairing for the wild type virus compared to N276D mutant virus. A 5-fold change in IC50 was deemed to be a significantly different change in phenotype. An increase in sensitivity is denoted in orange and an increase in resistance is denoted in blue. B. CD4bs bNAb IC50 change in the global panel of HIV env reference strains when N276D mutation is inserted. C. CD4bs bNAb IC50 change with N276D mutation versus N276K mutation.

To further assess the observed differential effects of N276D mutations on VRC01-class bNAbs sensitivity, we performed homology modeling of 12 HIV Env-VRC01 complex structures. A total of 12 HIV Env-VRC01 complex structures were modeled using Env sequences from a global panel of 12 viruses, with the BG505-VRC01 structure (PDB: 6V8X) as a template. In all modeled complexes, the side chain of glycan 276 showed nearly identical orientations and formed similar interactions with VRC01. This structural uniformity suggested that homology modeling was not sufficient to capture the observed functional effects associated with glycan 276 and support the idea that dynamics or other factors such as charge or glycan conformational rearrangements may govern the variation in neutralization sensitivity that we observed.

To test if the difference in neutralization was a result of glycan loss or residue change, the 246.F3-N276K virus was tested with the expanded panel of CD4bs antibodies (Fig 4C). N6 neutralized both N276 mutant viruses better than wildtype while 3BNC117 neutralized both wildtype viruses better than the N276 mutants suggesting that the glycan 276 in this 246.F3 Env shields N6 recognition but is necessary for 3BNC117 recognition. Clonally related antibodies VRC01 and VRC07-523 neutralized the N276D virus better than wildtype but were less potent against wildtype than against the N276K virus, suggesting that the residue was more important than the glycosylation status.

To further investigate the role of glycosylation, we produced a subset of wildtype and mutant pseudoviruses in Expi293F cells, which tend to add more high-mannose glycans to HIV-1 Env as compared to HEK293T cells. There was a slight loss of sensitivity to VRC01 for viruses produced by the Expi293F cell line as compared to HEK293T cells that did result in the mutant 246.F3 N276D virus becoming 5-fold more sensitive to the antibody as compared to the wildtype (completely opposite from the N276K phenotype) (Fig 5A). In general, all viruses in the X2278 background became more resistant to neutralization by all antibodies. However, the X2278 N276D mutant virus did not become more resistant to two CD4bs antibodies: VRC07-523 and N6 (Fig. 5B). Thus, these two antibodies still potently neutralized the virus without the N276 glycan when produced in Expi293F cells. These data suggest that the CD4bs epitope is more shielded from neutralization when more high-mannose glycans are present (wildtype virus produced in Expi293F cells), but that this increased neutralization resistance is lost when the N276 glycan is removed from the Env. The only other bNAb with differential neutralization profiles against viruses generated by the two producer cells was MPER bNAb 10E8v4-5R+100cF, with Expi293F viruses being more resistant to neutralization than HEK293T-produced viruses in all cases. These data suggest that this antibody is affected by the differential glycosyslation of the two producer cells (Supp Fig 2).

**Figure 5.**
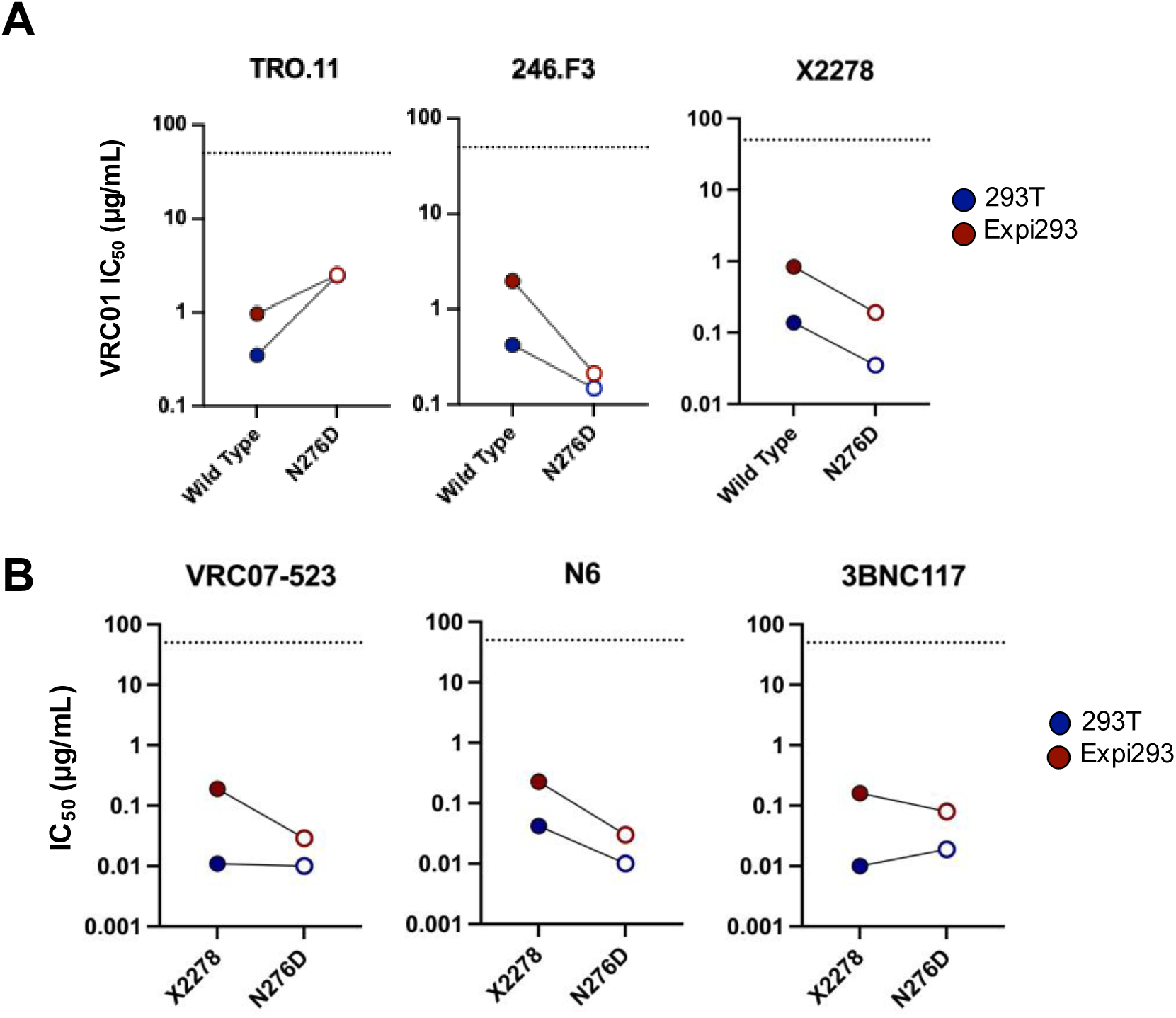
Effect of virus producer cell on neutralization sensitivity of viruses. **A**. Effect of producer cell differential glycosylation on VRC01 IC50 to 3 wildtype viruses and their matched N276D versions. B. IC50s of 3 different CD4bs antibodies to X2278 and X2278-N276D when produced by 2 different cell lines.

## Discussion

In this work, we found that removal of the N276 potential glycosylation site conferred varying levels of resistance to VRC01-class CD4bs bNAbs including VRC01, VRC07-523 and 3BNC117. In our *in vitro* virus escape assay, we determined that the escape mutation N276K in Env 246.F3 conferred complete neutralization resistance to VRC01. As N276 is the first residue of a potential N-linked glycosylation site, at first it was puzzling that its removal conferred resistance to VRC01 ^16,19,47^, as the removal of this glycan has been key to increasing the accessibility of naïve B cells to the CD4bs in vaccine studies ^14,15^. Despite this glycan being well-conserved, with 98% of analyzed sequences having N at this position (Fig 3B), and the ability to neutralize viruses with this glycan is necessary for antibody neutralization breadth, its removal should increase accessibility of the epitope. For some non-VRC01-like CD4bs antibodies for whom this glycan is a required part of the epitope, its removal is well documented to confer resistance, but this is not true for VRC01-class antibodies ^29^. Finally, virus circulating in the participant from whom VRC01 was isolated, added this glycan during co-evolution as a resistance mutation to the VRC01 lineage ^21^. For all these reasons, the loss of 276 glycan should not have conferred resistance to VRC01 neutralization. There are, however, examples of loss of the glycan inducing resistance. One study of soft randomization in the CD4bs of Env ADA (subtype B) found that changes to N276 that abrogated the putative N-linked glycan were associated with resistance to NIH45-46, 3BNC117 and VRC07 in their in vitro replication assay ^30^. In an *in silico* analysis of viral signatures of bNAb sensitivity, the presence of a glycan at position 276 is a sensitivity mutation for 3BNC117, suggesting that the removal of this glycan should confer resistance ^31^. Antigenic profiling, a method in which each amino acid change is made at all residues in an Env protein and incubated with bNAb to determine surviving populations, was performed using a subtype A Env BG505 library. In this assay, VRC01 did not select for the N276 glycan as an escape mutations, but a small number of viruses with the N276S mutation survived 3BNC117 selection, highlighting glycan removal could confer neutralization resistance to this bNAb ^32^. Importantly, these studies, in addition to ours, study virus escape *in vitro*, with these mutations arising only in the context of virus replicating in the presence of a bNAb, and therefore, other exogenous pressures, such as autologous antibodies, are not present. The lack of other host pressures could explain why removing this glycan is not commonly observed in circulating HIV Env sequences.

We studied the N276D mutation in the context of a global panel of HIV-1 Env reference strains and observed heterogeneity in the effects of the mutation on CD4bs bNAbs susceptibility, even those that are VRC01-class ^46^. Unsurprisingly, overall, there was a significant increase in the sensitivity of N276D viruses to CD4bs bNAbs VRC01, VRC07-523 and N6 as compared to wildtype. Interestingly, we observed more heterogeneity in the neutralization profile of N276D mutants against 3BNC117, with multiple viral strains, including 246.F3 (A1C), 25710 (C) and CH119 (CRF07_BC), gaining complete neutralization resistance, which matches the findings that 276 is a sensitivity mutation for this antibody. It is also worth noting that there was heterogeneity in which viruses became resistant to certain CD4bs bNAbs, highlighting that the same virus did not become more resistant to multiple CD4bs bNAbs. Some of this variability may derive from the fact that N276 in Loop D contacts the antibody light chain ^10^, and there is variability in the light chains of these VRC01 class antibodies. We did include control antibodies in our bNAb panel, non-CD4bs antibodies for which no change in neutralization sensitivity was expected, but the N276D mutations did confer a statistically significant increase in virus sensitivity to PGDM1400, which targets the V2 apex. It is possible that conformational changes induced by the glycan removal affected the binding epitope of PGDM1400 but also it is of note that while the sensitivity of the panel trended downward, only one circumstance conferred a greater than 5-fold change increase in sensitivity (Sup Fig 1A, 1B).

Finally, we observed that some of the changes in bNAb neutralization sensitivity may be due to amino acid changes and not associated with the loss of a potential glycosylation site. Our data comparing the N276D mutation to N276K suggest that 3BNC117 neutralization is more reliant upon the glycan at this position, whereas for VRC01 and VRC07-523, were sensitive to which residue was present. Of note, the change in 3BNC117 sensitivity was not equal between the N276D and N276K mutant viruses, suggesting that the loss of glycan but also the residue affected virus recognition. These differences could be explained because the mutation from an uncharged side chain (N) to either a positive charged side chain (K) or a negatively charged side chain (D) may cause structural perturbations that impact each antibodies’ recognition slightly differently. Finally, the complexity of the glycans modified by the producer cell line, can also affect bNAb sensitivity and how the loss of glycans affects bNAb recognition. We observed that certain wildtype viruses, especially X2278, were more resistant to neutralization when complex glycans were present (produced in Expi293F cells), but that this increased resistance could be lost when the N276 glycan was removed (i.e., N276D mutant virus produced in Expi293F cells remained potently neutralized).

Future studies should examine the role of different 276 residues in the context of global viruses and the impact on different bNAbs not only functionally but also structurally. These findings emphasize the importance of studying viral escape with a genetically diverse library to develop a deeper understanding of various escape pathways to better inform bNAb combinations as well as a deeper understanding of the influence of complex glycans.

## Acknowledgements

We acknowledge all of the individuals living with HIV from whom these viruses were isolated and make this work possible. The Env plasmids were obtained through the NIH HIV Reagent Program, Division of AIDS, NIAID, NIH: Panel of Global Human Immunodeficiency Virus Type 1 (HIV-1) Env Clones, HRP-12670, contributed by Dr. David Montefiori, and in particular the 246.F3 plasmid (ARP-12658) was contributed by Drs. Ronald Swanstrom, Li-Hua Ping, Jeffrey Anderson, David Montefiori, CHAVI, UNC-CH and Duke University. The antibody plasmids for heavy and light chains were obtained from the Vaccine Research Center (VRC). This research was funded in part by the National Institute of Allergy and Infectious Diseases of the National Institutes of Health grant number R01AG123456 to JB and RML and U01 AI169767 to KJB and RML. JN and QW were supported in part by NIH grant AI162236 and by UAB Research Acceleration Funds.

---

**Supplementary Figure 1.**
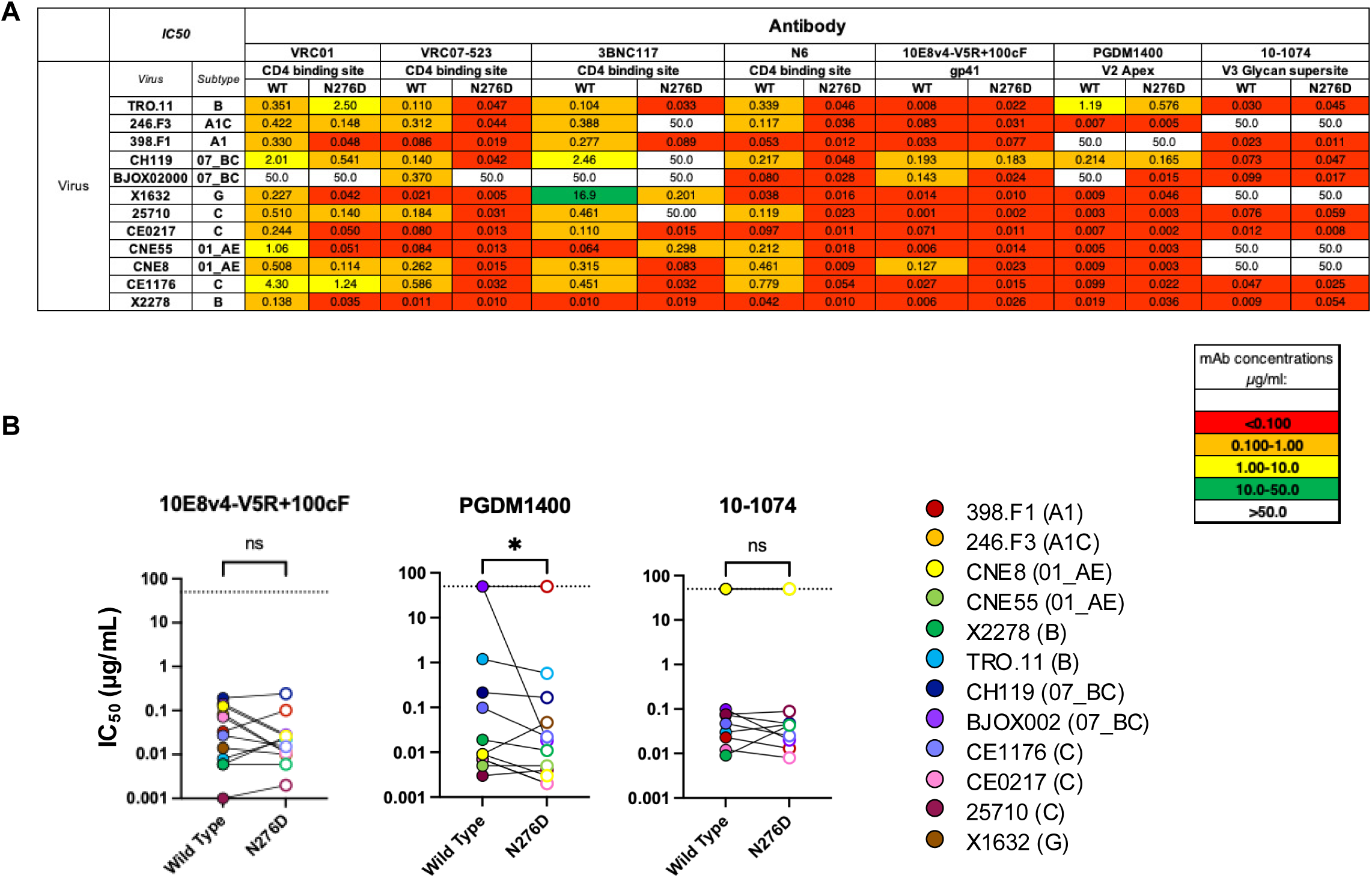
Effect of N276 glycan removal on neutralization profiles of bNAb panel. A. Heat map of the IC50 of each virus-antibody pairing for the wild type virus compared to N276D mutant virus. B. non-CD4bs bNAb IC50 change in the global panel of HIV env reference strains when N276D mutation is inserted.

**Supplementary Figure 2.**
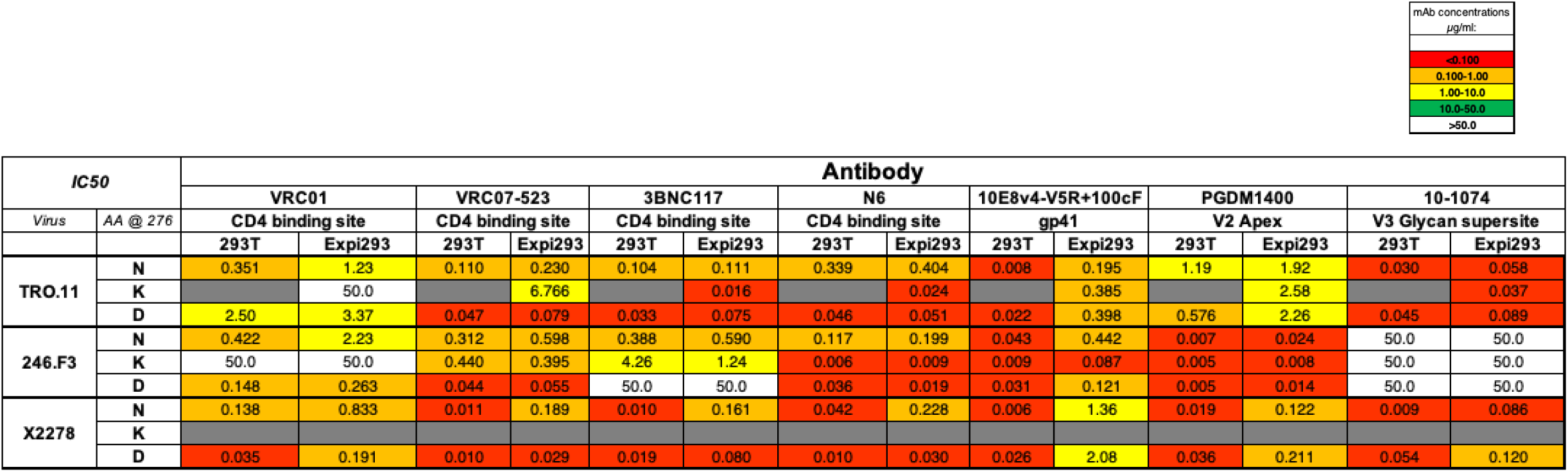
Effect of producer cell on mutant neutralization sensitivity. Heat map of the IC50 of TRO.11 and 246.F3 with each antibody pairing for the wild type virus compared to N276D and N276K mutant viruses. Gray indicates neutralization assays not performed.

